# Monitoring jumping ability and subjective recovery in women’s volleyball: acute effects of match play on countermovement jump and well-being

**DOI:** 10.1101/2025.07.08.663627

**Authors:** Arturo Franco, Pablo Casado, Manuel Conejero-Suárez, Jaime González-García

**Author notes:** Corresponding author: Jaime González-García, Universidad Francisco de Vitoria, Faculty of Health Science, *Ctra. Pozuelo-Majadahonda KM 1.800. 28223 Pozuelo de Alarcón, Madrid, España.

## Abstract

**Purpose:** Jumping ability is a crucial element for success in volleyball. Monitoring it helps optimize performance and prevent injuries. The countermovement jump (CMJ) is a reliable, non-invasive, and quick tool to periodically assess neuromuscular fatigue and performance, which, in combination with subjective measures, can offer a comprehensive view of fatigue/recovery status. The study aimed to: a) analyze changes in CMJ after a women’s volleyball match based on playing time and b) evaluate the temporal recovery of subjective variables TQR (perceived recovery) and HI (well-being). It was hypothesized that players would recover baseline values of CMJ and subjective variables 48 hours post-match.

**Methods:** A descriptive, longitudinal, and repeated-measures design was proposed. Twelve volleyball players were evaluated for changes in CMJ before and after a match, and at 24-, 48-, and 72-hours post-match. Outcome variables (jump height and RSImod), kinetic variables (concentric impulse), and jump strategy variables (contraction time and time to peak power) were analyzed. TQR and HI scales assessed perceived recovery and well-being.

**Results:** Significant changes were observed in jump height, concentric impulse, and RSImod between pre-match and 48 hours post-match in players with less playing time. Subjective variables TQR and HI did not show significant differences during the study period.

**Conclusion:** Competition did not significantly affect CMJ variables, suggesting that volleyball does not reduce jumping ability immediately after competing, which has direct implications for microcycle structure. The study highlights the importance of continuous monitoring of performance and recovery in volleyball.

## Introduction

Jumping ability is a crucial element for success in volleyball. The main actions performed by players during the game largely depend on their jumping ability (e.g., spike, block, serve…). Additionally, a relationship between jumping ability and the effectiveness of offensive actions has been demonstrated (1,2). Similarly, differences in jumping ability among different levels of competition have been observed in both men and women (3,4).

To maximize performance, it is necessary to monitor athletes with the highest possible frequency, which allows to identify the relationship between load and injury risk. This process involves the precise measurement and monitoring not only of the sports and non-sports loads that athletes face but also of their performance, emotional well-being, symptoms, and injuries (5). The benefits of monitoring athletes are numerous: it allows for explaining changes in performance, increasing the understanding of training responses, detecting fatigue and recovery needs, as well as informing the planning and modification of training programs and competition schedules. Additionally, it ensures the generation of appropriate load doses to minimize the risk of non-functional overtraining, injuries, and illnesses (6).

In addition to being related to performance, jumping ability is a sensitive marker of fatigue, which can help identify athletes’ readiness for the next competition and/or training session. This, in turn, assists coaches and physical trainers in managing the training load in subsequent sessions (7,8). Due to the aforementioned factors, its reliability, and the speed in collecting and analyzing data, the countermovement jump (CMJ) is widely used to assess athletes’ jumping ability and the effectiveness of different training programs (9). This assessment of lower-body neuromuscular performance has demonstrated excellent reliability both within and between days in volleyball players (8), making it a comprehensive tool for frequent use in performance and fatigue evaluation processes in ecological contexts. With an appropriate metric selection, CMJ can provide valuable information for athlete performance profiling, monitor neuromuscular fatigue, or tracking athletes’ recovery status after injury (7). Given its characteristics, the analysis of alternative metrics to traditional ones is an excellent method for monitoring athlete fatigue. It provides coaches with valuable insights to ensure proper neuromuscular recovery and optimize decision-making in training, thereby enhancing the effectiveness of training adaptations. However, given the multidisciplinary and holistic nature of fatigue etiology, using markers like the CMJ alone does not provide a complete view of the process of training stimulus, fatigue, response, and adaptation. Therefore, it is advisable to include subjective assessments of their well-being, in addition to monitoring training and competition loads and evaluating neuromuscular fatigue. These assessments provide complementary information to the neuromuscular fatigue markers derived from evaluations like the CMJ (10). Total Quality Recovery Scale (TQR) (11) and Hooper Index (HI) (12), are two subjective evaluations of athlete’s well-being that complement neuromuscular fatigue analysis. TQR evaluates from 1-10 the perceived recovery status of athletes using a visual analogic scale. It correlates with heart rate variability parameters in female volleyball players (13), and has also been shown as sensible to weekly training load variations in volleyball (14). Hooper Index (HI) and its sub-items (sleep quality, fatigue, stress, and muscle soreness) are also a promising tool for monitoring fatigue in team sports (15). It has shown an association with training load in professional football (16), with reduced values observed up to 72 hours post-match. To optimize adaptation and recovery, it is necessary to use objective methods for monitoring loads, fatigue, and recovery, given its holistic nature. This should be combined with subjective assessments of recovery and form. However, there is currently no information that has analyzed the neuromuscular recovery profile through the analysis of unidimensional metrics derived from the CMJ in conjunction with subjective scales of well-being and recovery in relation to training and competition load. Therefore, the aim of this study was to monitor the recovery profile of jumping ability and perceived well-being of the players based on metrics derived from CMJ and TQR according to match time.

## Methods

### Design

To address the research objectives, a descriptive, repeated measures, observational, multivariable, longitudinal, and prospective design was employed. During the first two sessions, participants were familiarized with the CMJ test, and the reproducibility and sensitivity of its metrics was established. The following four observations constituted the experimental period of the study. In Observation 1 (match day, MD), the CMJ was performed before the official warm-up. Observations 2, 3, and 4 were conducted immediately post-match, at 24, 48, and 72 hours (Figure 1), respectively. All observations were carried out after the warm-up.

**Figure 1.**
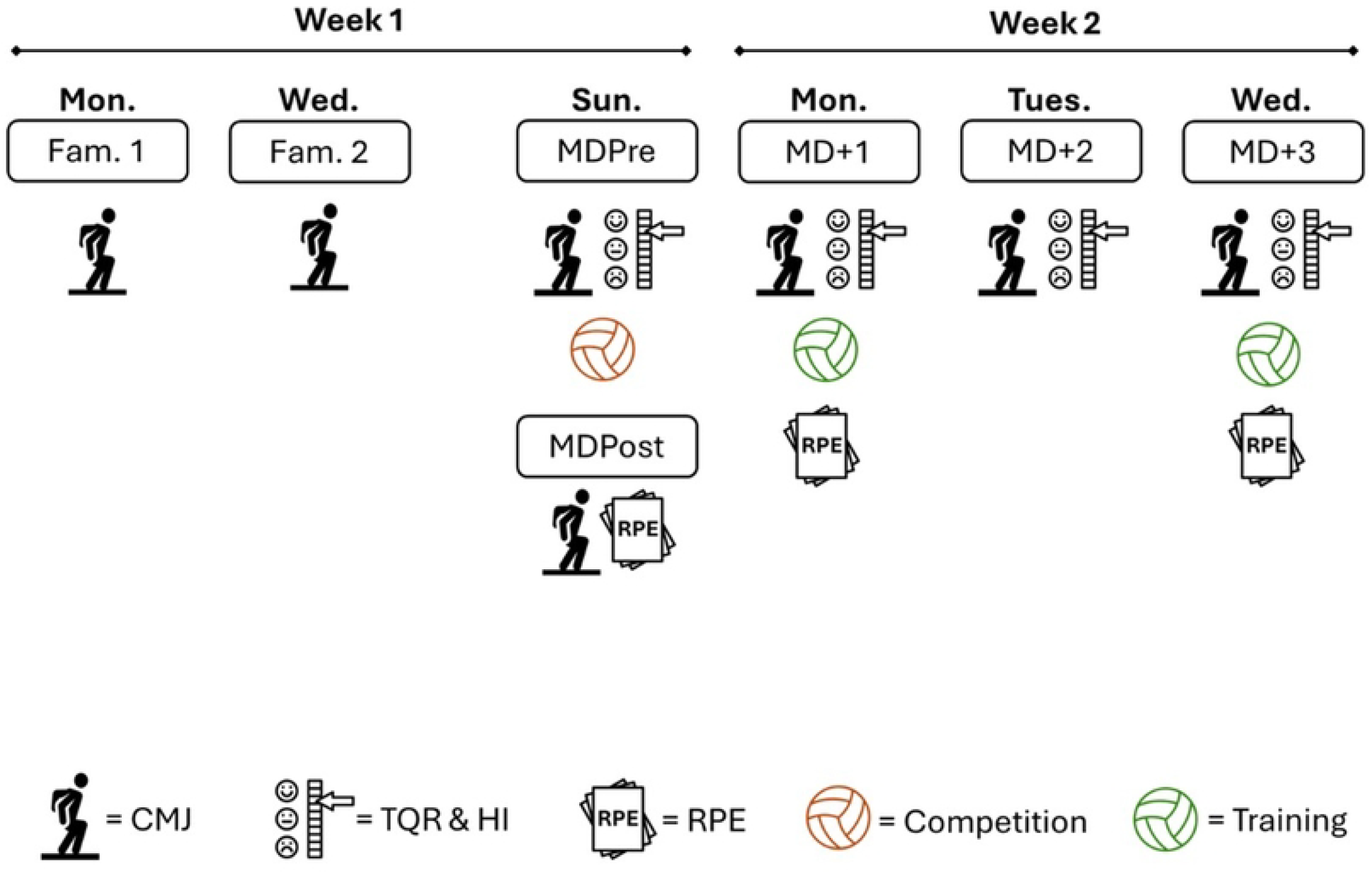
Experimental design and assessment timeline for analyzing post-match recovery profiles in starting and non-starting volleyball players.

### Participants

Twelve female volleyball players were involved in the study (22.8±3.39 years; 168±6.01 cm in height; 63.9±4.68 Kg in weight; 6±3 years of competitive experience). They are competitors in the second division of the Madrid Volleyball Federation during the 2023/2024 season. To be included in the study, participants must have had at least 2 years of volleyball training experience, have not suffered any injuries in the past six months. The data were collected prospectively from February 1st to February 15th, 2025. All participants provided written informed consent prior to participation. The research project was approved by the Ethics Committee of the ***blinded for reviewing purposes*** and is registered under number 10/2024. The study complies with the requirements of the Helsinki Declaration.

### Warm-up

Prior to each assessment, a standardized warm-up following the RAMP Model (Rise – Activate and Mobilize – Potentiate) was conducted. This warm-up included 5 minutes of jogging, 5 minutes of dynamic stretching, mobility exercises (for the shoulders and hips), and core activation, finishing with 1 set of 5 CMJs and 1 set of 5 CMJs with rebound. Rest intervals of 2 minutes were allowed between each warm-up block. To ensure optimal performance in the subsequent evaluations, the intensity of the warm-up progressively increased.

### CMJ evaluation

A dual force platform Force-DecksFD4000 system (ForceDecks, London, United Kingdom) was used to evaluate CMJ. The platforms were calibrated beforehand following the manufacturer’s instructions. They recorded at a frequency of 1000 Hz. Participants performed 3 attempts, with 10-15 seconds of rest between each. They were instructed with the verbal command: “hands on hips, maximum speed, and maximum height.” Before the first jump, participants were asked to stand on the platforms and remain stationary for 1 second until the platforms completed the weight measurement. The start of the jump was defined as the moment when a reduction in vertical ground reaction force of at least 20 N below the subject’s body weight was detected. Body weight was determined during a weighing phase of at least one second of quiet standing.

The following variables were analyzed using a forward dynamics approach. Jump outcomes included jump height, calculated from the force-time data using the impulse-momentum relationship, and the modified Reactive Strength Index (RSImod), obtained by dividing jump height by contraction time. Kinetic variables comprised concentric impulse, computed as the integral of vertical ground reaction force over time during the propulsive phase. Jump strategy metrics included contraction time, defined as the interval from movement onset to take-off, and time to peak power.

### Training and competition loads quantification

To analyze the match demands, an arbitrary units measure will be used, calculated from the equation “Load = RPEsession*Session Duration (minutes)” (17). This method has shown strong associations with heart rate data or global positioning systems (GPS) in team sports (17). RPE was reported by participants 10 minutes after the match or training session ended (17). To ensure that the training load of the MD+1 session is consistent across all players, the same formula was used. Descriptive data of training load is shown in Table 1.

**Table 1.** Differences in training/competition load between groups.

| Play-time group | TL MD (Au) | TL MD+1 (Au) |
| --- | --- | --- |
| High | 238±127.86 | 234±186.62 |
| Low | 46.4±60.02† p=0.011 | 450±269.95* p=0.048 |

*Note.* TL = training load; †=significant with respect to

### Total Quality Recovery and Hooper Index

To evaluate perceived recovery status, the Total Quality Recovery (TQR) scale was used, which employs a Likert scale from 1 to 10, with 10 indicating the highest possible perceived recovery. For assessing well-being, the Hooper Index (HI) was utilized, which evaluates sleep quality, fatigue, muscle soreness, and stress on a scale from 1 to 7 each. The Hooper Index is calculated by summing the scores of the 4 questionnaire items. This value can range from 4 to 28, with a lower score indicating better well-being. Both assessments were administered via a Google Forms questionnaire sent out every morning (9:00–10:00 am).

### Statistical Analysis

For statistical analysis, Jamovi software (Jamovi) was used. The normality of the data distribution was assessed using the Shapiro-Wilk test. Intraclass correlation coefficients (ICCs) (18) with 95% confidence intervals (CI 95%) were evaluated using the following criteria: poor reliability (<0.5), moderate reliability (0.5-0.75), good reliability (0.75-0.90), and excellent reliability (>0.90) (19). Grouping by playing time (high or low) was performed using median split analysis (high play time group = >40 min; low play time group = ≤40 min.). A two-way ANOVA was conducted to identify differences in recovery profiles between players with high and low play time. Bonferroni correction was applied for post hoc comparisons to reduce the risk of Type I errors associated with multiple testing.. Effect sizes (Cohen’s d) between conditions were calculated and classified as follows: ≤0.2 (trivial), ≥0.2-0.6 (small), ≥0.6-1.2 (moderate), ≥1.2-2.0 (large), and ≥2 (very large) (20). Statistical significance was set at p<0.05. Results are presented as mean ± standard deviation.

## Results

The variables jump height (ICC = 0.98 [0.01-0.99]; SEM = 0.66 cm; MDC 95% CI = 1.83 cm; Mean CV = 1.83%) and concentric impulse (ICC = 0.99 [0.94-1.00]; SEM = 2.05 N/s; MDC 95% CI = 6.68 N/s; Mean CV = 1.11%) demonstrated excellent reliability. The modified reactive strength index (RSI) (m/s) (ICC = 0.78 [0.32-0.94]; SEM = 0.04 m/s; MDC 95% CI = 0.11 m/s; Mean CV = 6.72%) showed good reliability. In contrast, contraction time (ICC = 0.45 [-0.23-0.84]; SEM = 63.41 ms; MDC 95% CI = 175.76 ms; Mean CV = 6.03%) and time to peak power (ICC = 0.46 [-0.24-0.84]; SEM = 0.06 s; MDC 95% CI = 0.18 s; Mean CV = 6.61%) exhibited poor reliability.

A main effect of time was observed for the variables jump height [F(1.557, 14.01) = 7.006; p=0.0111] and modified reactive strength index (RSI) [F(1.621, 14.59) = 3.964; p=0.0018]. No significant effect of the group based on playing time was found, nor was there a significant interaction effect for any variable.

**Figure 2.**
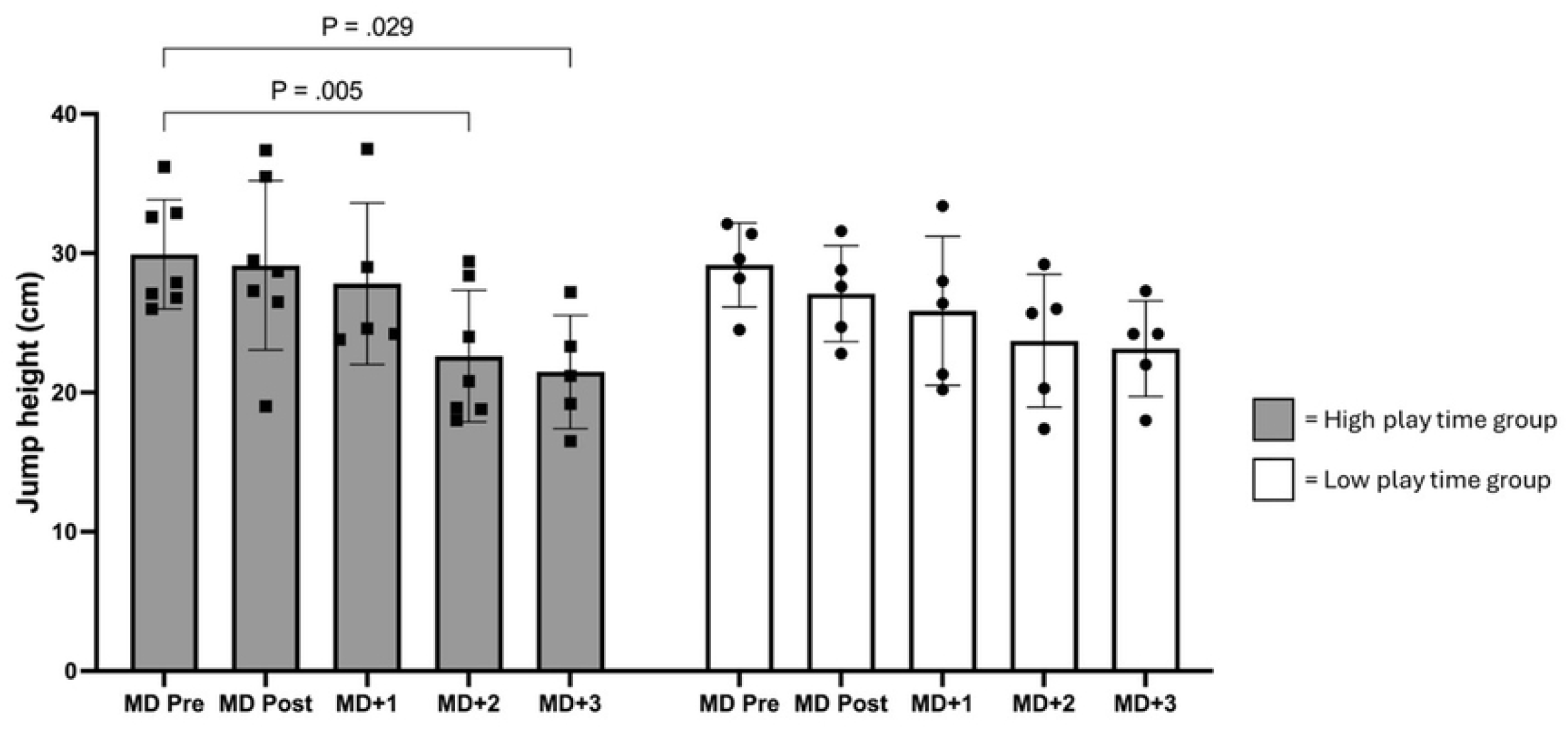
Changes in countermovement jump (CMJ) height across time points in high and low play time volleyball players.

**Figure 3.**
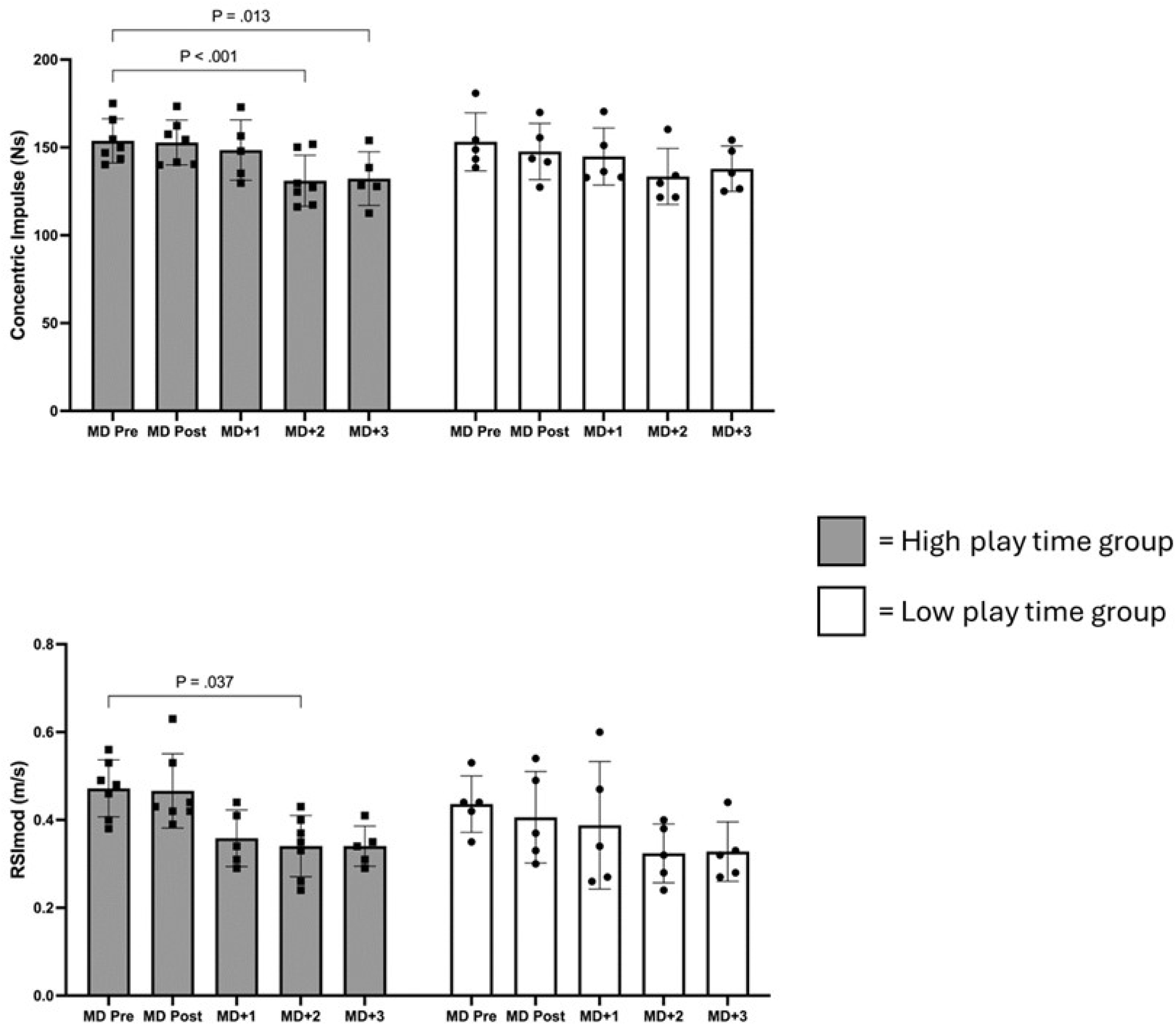
Evolution of concentric impulse and RSImod following match play in high and low play time volleyball players.

In the high playing time group, a reduction in jump height (Figure 2), concentric impulse, and modified RSI was observed at MD+2 compared to MDPre [Jump height: ES = -1.62 [-2.10 to - 1.25]; p=0.005; Concentric impulse: ES = -1.57 [-1.78 to -1.37]; p<0.0001; RSImod: ES = -1.57 [-2.31 to -1.21]; p=0.004], and at MD+3 compared to MDPre [Jump height: ES = -2.08 [-2.47 to -1.70]; p=0.029; Concentric impulse: ES = -1.61 [-1.87 to -1.36]; p=0.001; RSI mod: ES = -1.04 [-2.72 to -1.35]; p=0.025] (Figure 3). In the low playing time group, significant differences were observed only between MDPre and MD+2 in concentric impulse (ES = -0.6 [-2.67 to -1.48]; p=0.034) and modified RSI (ES = -0.6 [-2.67 to -1.48]; p=0.022). For contraction time and time to peak power, no influence of the measurement time was observed in either group (Figure 4).

**Figure 4.**
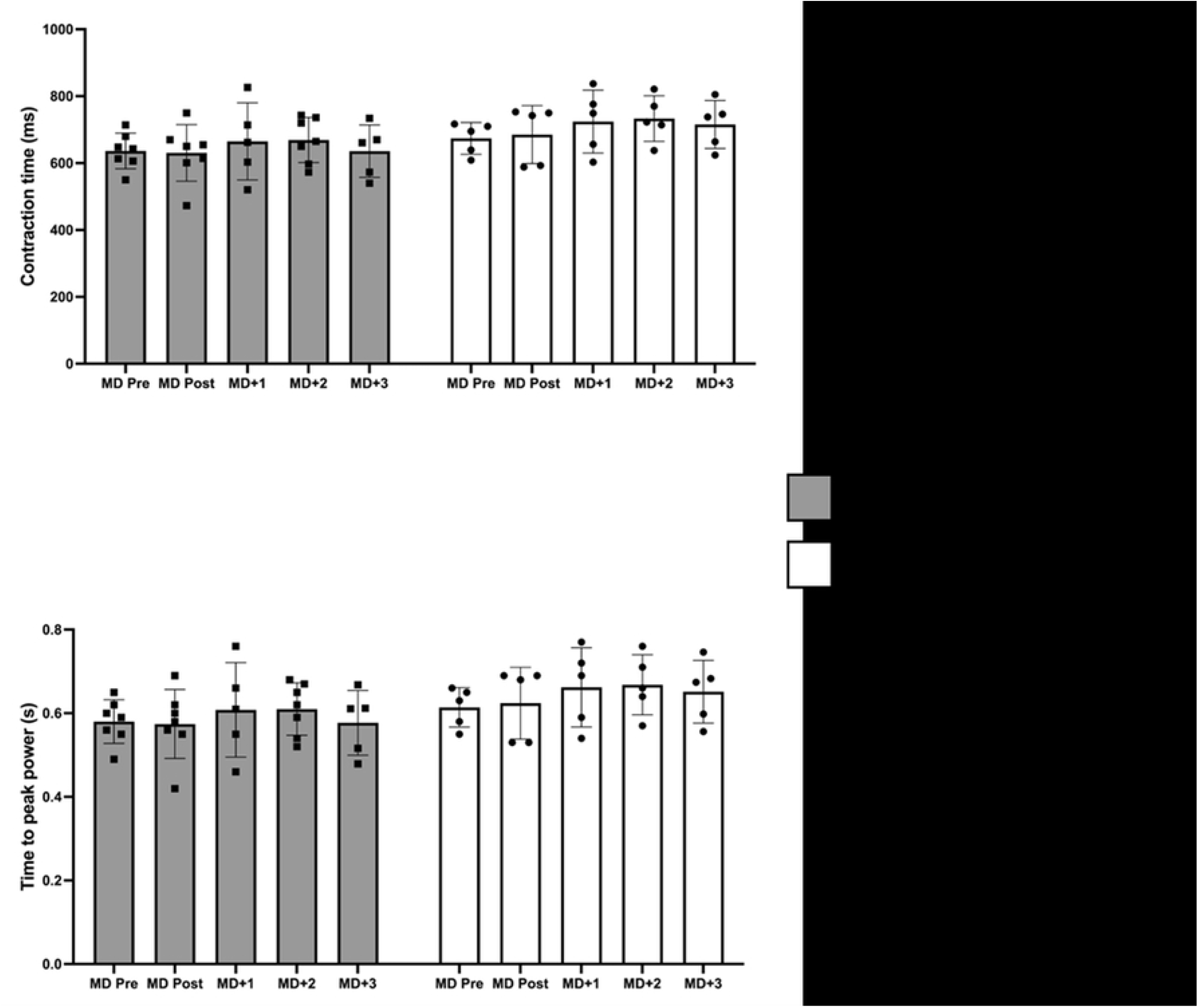
Contraction time and time to peak power in high and low play time volleyball players across the post-match recovery period.

For the subjective variables TQR and HI, there were no significant differences between match day and MD+1, MD+2, and MD+3 in either group (p>0.05) (Figure 5).

**Figure 5.**
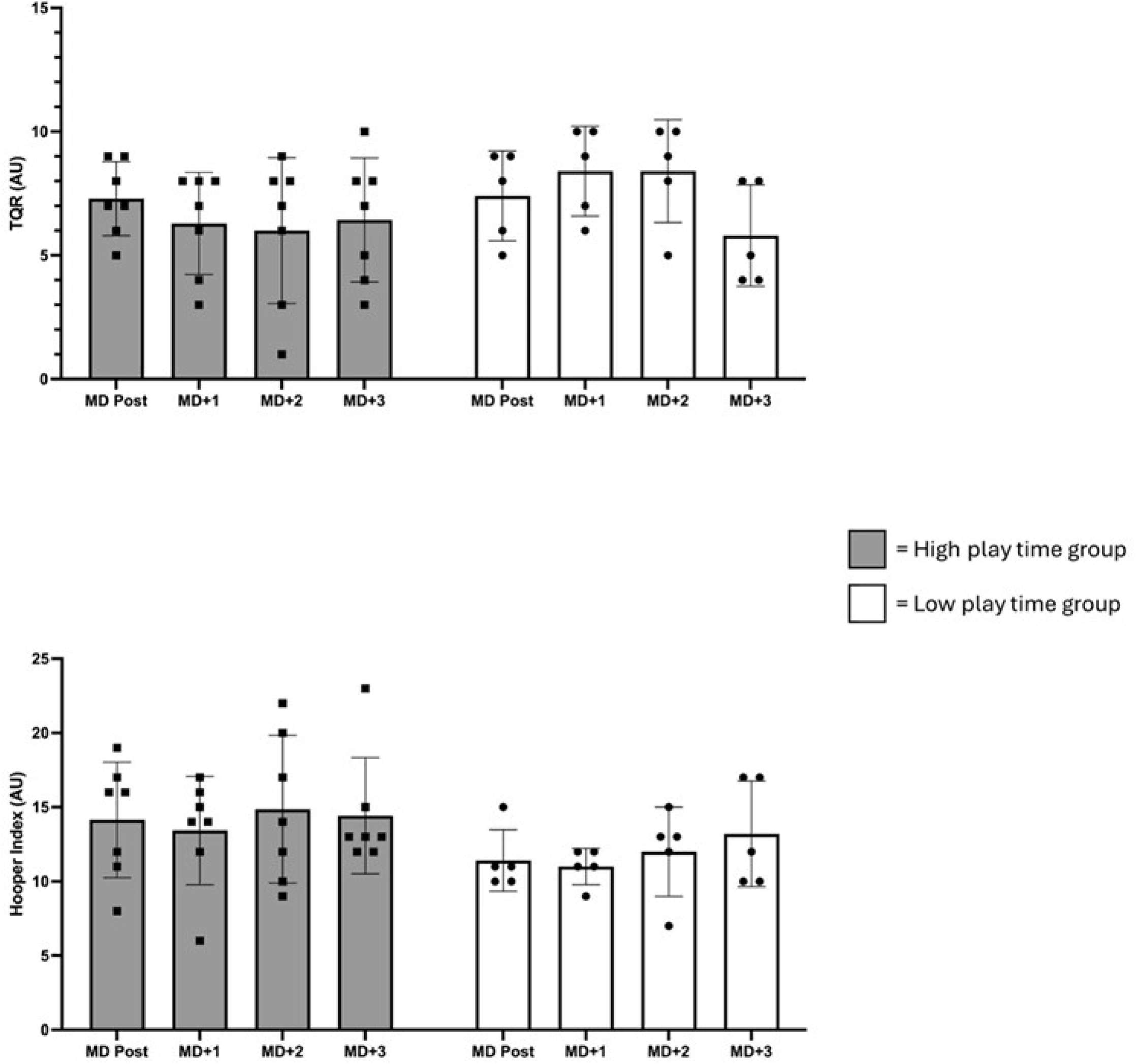
Perceived recovery (TQR) and wellness (Hooper Index) following match play in high and low play time volleyball players.

## Discussion

The primary objective of the study was to identify the recovery profile of various CMJ metrics and perceived recovery following a semiprofessional women’s volleyball match, based on play time. Our findings indicate that the match (load Au range: 46.4 – 238.9) did not influence jump outcomes (jump height and modified RSI), kinetics (concentric impulse) and strategy metrics (contraction time and time to peak power). However, significant changes were detected in jump height, concentric impulse, and modified RSI (Outcomes and kinetics) between MDPre and MD+2. No significant changes were observed for TQR and HI in any of the assessments. Neuromuscular capabilities changes can be explained by the influence of the MD+1 training session, which involved a higher load compared to the match (Table 1), especially for players in the low play time group.

Our results indicate that the match did not influence any of the CMJ variables, regardless of play time group. Previous research has observed the same recovery pattern in professional male volleyball players (21), with no changes in jump height and contraction time (jump strategy) after across two matches. Other studies aimed to measure changes in jump height throughout microcycles using inertial systems (22). In these studies, no changes were observed in mean and maximum jump height were observed regardless of the microcycle session and time until next competition. All these findings demonstrate that the physiological demands of volleyball competition do not lead to a reduction in athletes’ jumping ability immediately after competing, which would have direct implications for the structure of microcycles, allowing for high-load on the following sessions. In contrast, in other team sports, competition has shown greater reductions in post-match jump capacity, kinetics, and kinematics. In fact, reductions of between 1.6 and 6 cm have been observed (23). This can be explained by the differences in the physiological demands of the sports. In volleyball, the duration of each point ranges from 3 to 40 seconds, followed by a brief recovery period averaging 12 seconds (24). Additionally, the duration of the competition is typically shorter, although it depends on the progression of the match (66 minutes in the game analyzed). These differences in physiological demands (volume and density), combined with the high specialization of volleyball players in actions involving the stretch-shortening cycle (25),may explain the variation between volleyball and other sports in terms of reductions in jump height and the strategies employed by athletes, and consequently, in the fatigue generated by the competition.

According to our results, jump height remained stable until MD+2. Given the recovered jump capacity at 24 hours post-competition, the microcycle periodization model allows for the introduction of an impact training session on MD+1 (Table 1). This session (training load shown in Table 1) did lead to a decrease in jump results (height ES [95%CI]: -0.74 [-1.52 to 0.04] and RSI mod ES [95%CI]: -0.45 [-1.01 to 0.10]) and kinetics (concentric impulse ES[95%CI: -0.92 [-1.57 to -0.27]) but did not affect jump strategy (contraction time ES[95%CI]: -0.04 [0.32 to - 0.25] and time to peak power ES[95%CI]: -0.04 [-0.33 to 0.25]), while TQR (ES[95%CI]: -0.28 [-1.08 to 0.51]) and HI (ES[95%CI]: 0.27 [-0.37 to 0.9]) scales presented small changes. This contradicts the findings of Bishop et al., (2023), which suggest that fatigue is observed through changes in jump strategy. The high specificity in adaptations due to ballistic training (fast contractions) and the long intrinsic recovery periods in volleyball may prevent rapid force production from being affected, allowing for greater recruitment of muscle fibers, resulting in minimal changes in CMJ metrics (26). In these terms our results indicate that the most fatigue-sensitive metrics in volleyball are jump height, concentric impulse, and RSI mod.

Following competition in both male (27) and female soccer (28), and after a training volume similar to that of the MD+1 session (90 minutes), reductions have been observed in the capacity to develop maximum power in a CMJ, jump height, and other physiological variables, which may not fully recover until after 48 hours and could require up to 72 hours to return to baseline levels (27). In the case of rugby, the same phenomenon has been observed (23). Our results, along with those observed previously, suggest that there is a dose-response relationship between load and the subsequent recovery periods. Given all this information, it is essential to monitor the post-training response according to the training load to identify the recovery periods. This approach helps to minimize the risk of overtraining and maximizes neuromuscular adaptations to the training program (5). Following our results and the results obtained in other sports, it seems necessary to allow 24 to 48 hours between a fatiguing training or competition stimulus and the next session, if the goal is to arrive with minimal fatigue for the subsequent session.

Based on previously observed recovery periods in team sports, it has been suggested that the session following a match should focus on recovery and involve a reduced training volume (22). Our results suggest that in a context where the match imposes less stress than the training itself, it may not be essential to conduct a recovery session after the competition. This is because athletes have already regained their neuromuscular capacity without affecting TQR and HI values. Identifying individual recovery periods for each context or club will enable coaches to schedule high-intensity training sessions the day after a competition. This approach can lead to greater chronic load accumulation and progress training loads with a reduced risk of injury (29). Higher chronic load values are associated with a lower risk of injury, as well as improved aerobic and athletic performance (30). Coaches and strength and conditioning professionals should analyze their context and design microcycles utilizing periods of lower fatigue to apply training load, especially in amateur settings where weekly training hours are limited. In the case of volleyball, it may be advisable to conduct an intense training session on MD+1, provided that the state of recovery allows it. To effectively monitor neuromuscular capabilities in volleyball, it results more reliable and sensitive (8) to use force time derived metrics over other systems used previously such as inertial sensors. We recommend the use of this method to optimize the comprehension and decision making in terms of monitoring fatigue and structuring microcycles.

This study presents several limitations that should be acknowledged. First, the relatively small sample size may limit the generalizability of the findings. Second, the absence of a control group prevents causal inferences and limits the ability to compare the observed outcomes against a standardized reference. Lastly, data were collected from a single match, which restricts the representativeness of the results and may not capture variability across different competitive contexts or time points.

## Conclusion

In conclusion, a semi-professional women’s volleyball match does not induce changes in jump outcomes, kinetics, or jump strategy regardless of playing time. This supports conducting high-intensity training on MD+1, allowing for greater load accumulation. Coaches and strength and conditioning professionals should use a combination of neuromuscular tests (CMJ) and subjective athlete assessments (TQR and HI), as relying on either approach in isolation may lead to errors in training decision-making.

